# Premorbid performances determine the deleterious effects of nigrostriatal degeneration and pramipexole on behavioural flexibility

**DOI:** 10.1101/2022.08.24.505066

**Authors:** Mélina Decourt, Eric Balado, Maureen Francheteau, Marcello Solinas, Marianne Benoît-Marand, Pierre-Olivier Fernagut

## Abstract

Subtle cognitive impairment can occur early in the course of Parkinson’s disease (PD) and may manifest under different forms of executive dysfunction such as impaired cognitive flexibility. The precise contribution of nigrostriatal dopaminergic neurodegeneration to these non-motor features of the disease is poorly known. Whether such cognitive impairment associated with the disease process may also predate and contribute to the development of neuropsychiatric side-effects following dopamine replacement therapy remains largely unknown. To address these issues, we investigated the respective contributions of nigrostriatal degeneration and chronic treatment with the dopamine D3-preferring agonist pramipexole on behavioural flexibility in a rat model of PD. Flexible, intermediate and inflexible rats were identified based on baseline assessment of behavioural flexibility using an operant set-shifting task. Nigrostriatal degeneration was induced by bilateral viral-mediated expression of A53T mutated human _α_-synuclein in the substantia nigra pars compacta and behavioural flexibility was assessed after induction of nigrostriatal degeneration, and during chronic pramipexole treatment. Nigrostriatal degeneration impaired behavioural flexibility in flexible but not in inflexible rats. Pramipexole induced a decrease of behavioural flexibility that was exacerbated in lesioned rats and in the most flexible individuals. Furthermore, the deficits induced by pramipexole in lesioned rats affected different components of the task between flexible and inflexible individuals. This study demonstrates that nigrostriatal degeneration and pramipexole unequally impair behavioural flexibility, suggesting that the susceptibility to develop non-motor impairments upon treatment initiation could primarily depend on premorbid differences in behavioural flexibility.

## INTRODUCTION

Non-motor features of Parkinson’s disease (PD) and non-motor side effects of dopamine replacement therapy are important factors contributing to the decreased quality of life of the patients.^1-3^ Within the large spectrum of non-motor symptoms, subtle cognitive impairment can occur early during the course of the disease in one third of patients and may manifest under different aspects of executive dysfunction, such as attentional, memory deficits, as well as decreased cognitive flexibility.^4-6^

Dopamine replacement therapy with dopaminergic D2/D3 agonists is widely used to relieve motor symptoms but can be associated with compulsive behaviours designed as impulse control disorders (ICD) in a significant proportion of patients.^7-10^ ICD are defined by the inability to refrain the urge to repeatedly engage in behaviours leading to harmful personal, social or financial consequences such as pathological gambling, hypersexuality, binge eating or compulsive shopping.^10^ The fact that ICD affect only a subset of PD patients suggests a significant role for an underlying individual vulnerability as a predisposing factor.^3^

Executive functions contribute to develop adapted goal-directed behaviours and several clinical studies have highlighted impairments in PD patients with ICD.^11-13^ Particularly, working memory, visual-spatial planning, reward processing and cognitive as well as behavioural flexibility (set-shifting, task switching) are impaired.^11,13-15^ Cognitive flexibility represents the ability to switch thinking between concepts and behavioural flexibility is defined as the ability to adapt behaviour according to environmental changes. As tests assessing cognitive flexibility require behavioural responses, both constructs are intimately associated.^16^ Whether such type of cognitive impairment associated with the disease process may predate and/or contribute to the development of ICD remains largely unknown.

The pathophysiology of executive dysfunction involves dopaminergic and non-dopaminergic (cholinergic, noradrenergic and serotonergic) systems, leading to the dysfunction of frontostriatal circuits.^17^ Interestingly, frontostriatal dysfunction has been identified in PD patients with ICD ^12,18^, suggesting some pathophysiological overlap between non-motor symptoms of PD and non-motor side effects of dopamine replacement therapy. The occurrence of these non-motor features early in the course of the disease further suggest a possible contribution of nigral dopaminergic neurons.

Here, we used a rat model of PD induced by viral-mediated expression of _α_-synuclein in the substantia nigra pars compacta (SNc) ^19,20^ to probe the potential contribution of nigrostriatal degeneration to behavioural flexibility and its possible implications for developing further impairments under dopamine replacement therapy. To this end, we investigated interindividual responses to nigrostriatal degeneration on behavioural flexibility in a set-shifting task, as well as the effects of pramipexole (PPX), one of the most widely used dopaminergic treatment that is also implicated in ICD.

## RESULTS

### Characterization of the population based on basal inflexibility score

Upon completion of baseline sessions (**Fig 1A**), the analysis based on the inflexibility score allowed segregating three different groups (H = 45.50, df =2, p<0.0001, **Fig 1B**): FLEX, INT and INFLEX (FLEX vs INT: p<0.01, INT vs INFLEX: p<0.01, FLEX vs INFLEX p<0.0001). Baseline error rate was higher for the egocentric dimension than for the visual dimension in all groups (F_1, 100_ = 47.02, p<0.0001; FLEX: p<0.05, INT: p<0.001, INFLEX: p<0.001, **Fig 1C**) indicating better performances when the active lever was indicated by a visual cue. Moreover, INFLEX animals made more errors than FLEX animals both in the visual and egocentric dimensions (F_2, 100_ = 14.15, p<0.0001; visual: p<0.05, egocentric: p<0.001, **Fig 1C**).

**Figure 1:**
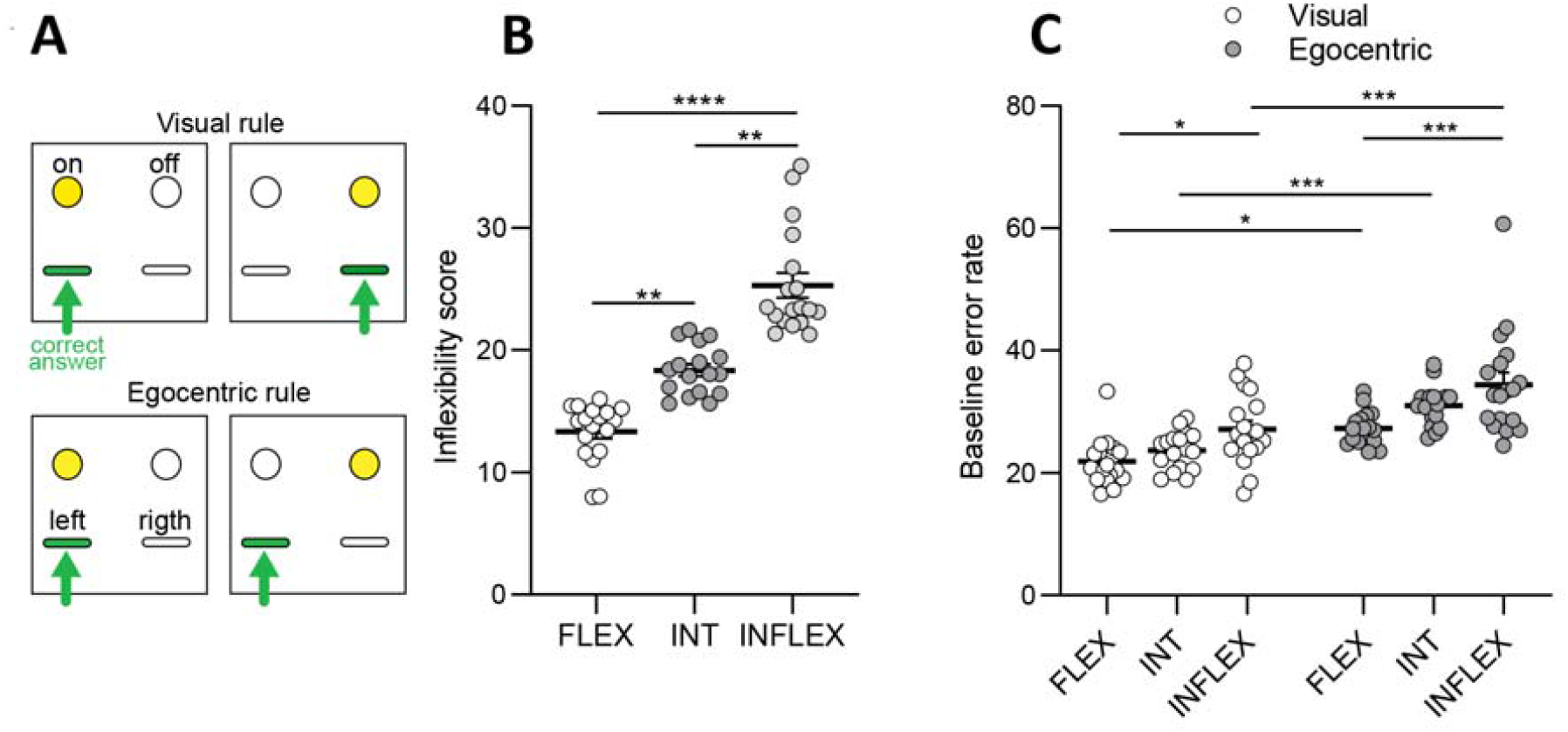
Overview of the set shifting task and baseline categorization of Flexible, Intermediate and Inflexible rats. **A**. Presentation of the set shifting task. In the visual rule, the position of the light cue indicates which lever is rewarded (shown in green). In the egocentric rule, one lever (either left or right) is rewarded and the position of the light cue is random. **B**. Inflexibility scores of Flexible (FLEX), Intermediate (INT) and Inflexible (INFLEX) rats at baseline. **C**. Baseline error rates in the visual and egocentric rules for FLEX, INT and INFLEX rats. *, p<0.05, **, p<0.01, ***, p<0.001.

### Alpha-synuclein-induced nigrostriatal degeneration and motor deficits

Stereological counts showed that viral-mediated expression of _α_-synuclein led to a significant loss of TH-positive neurons in the SNc (-42.56%, t = 9.898; df = 51; p<0.0001 **Fig 2A**). The extent of the lesion was independent of baseline flexibility and each group that underwent viral-mediated expression of _α_-synuclein presented a similar loss of SNc neurons compared to their sham counterparts (**Fig 2B**).Viral-mediated expression of _α_-synuclein in the SNc induced progressive bilateral motor deficits with a significant effect of the lesion (F_1, 204_ = 316.8, p<0.0001), time (F_3, 204_ = 77.04, p<0.0001) and lesion X time interaction (F_3, 204_ = 45.23, p<0.0001, **Fig 2C**). Motor deficits were significant at 4 weeks post-AAV (p<0.0001), worsened at 9 weeks (p<0.0001) and were improved by PPX (p<0.0001). The magnitude of stepping deficits was similar between all lesioned rats regardless of the initial inflexibility score (**Fig 2D**).

**Figure 2:**
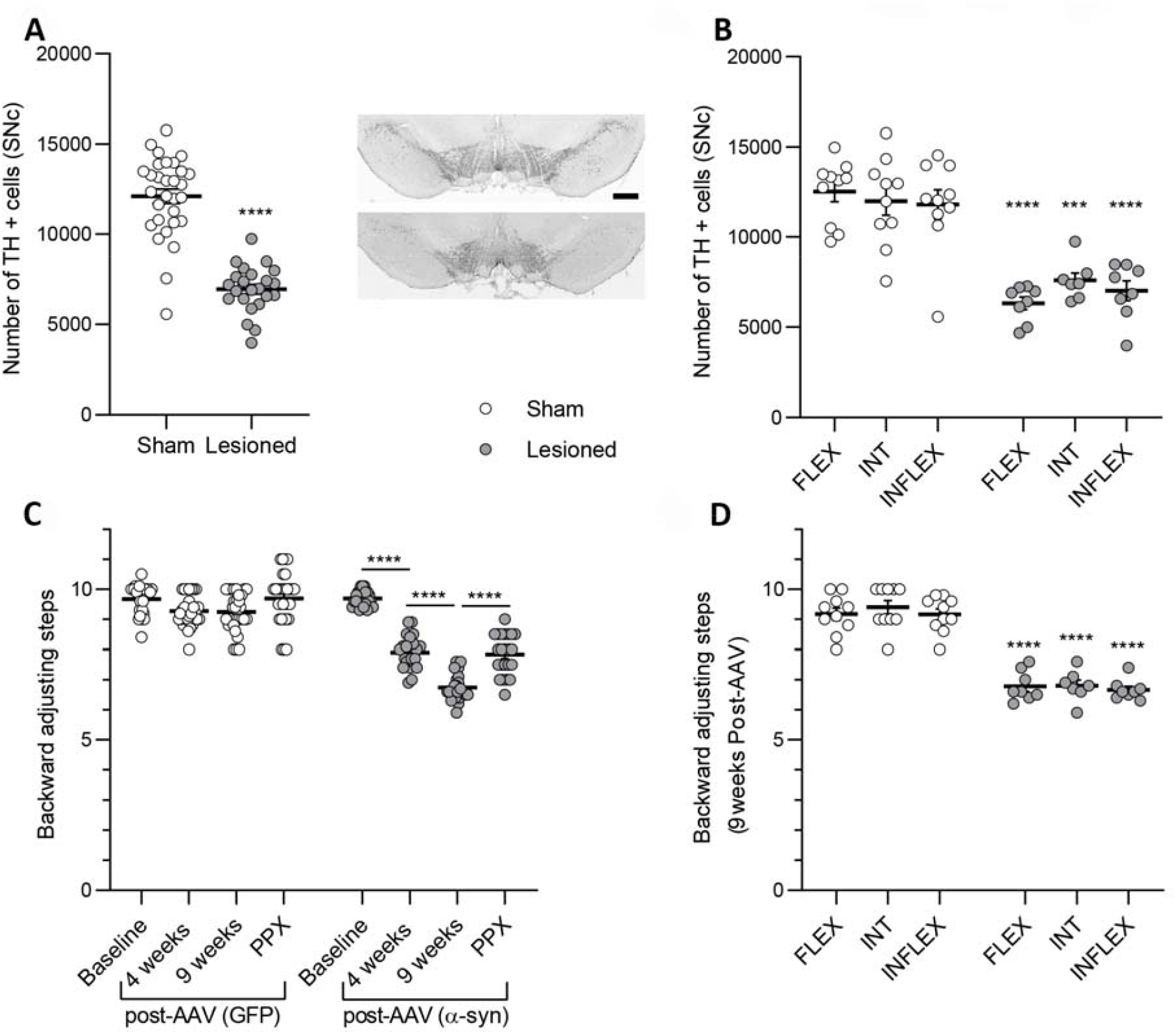
alpha-synuclein induced similar neurodegeneration and motor deficits regardless of baseline behavioural inflexibility. **A**. Stereological counts of tyrosine hydroxylase (TH) positive neurons in the SNc and representative images of the SNc in sham (top) and lesioned rats (bottom). Scale bar = 1 mm. **B**. Magnitude of neurodegeneration in FLEX, INT and INFLEX rats. **C**. Motor impairments induced by viral mediated expression of alpha-synuclein in the SNc at 4 and 9 weeks post-AAV injection are alleviated by pramipexole (PPX). **D**. Magnitude of motor impairments at 9 weeks post-AAV injection in FLEX, INT and INFLEX rats. *, p<0.05, **, p<0.01, ***, p<0.001, ****, p<0.0001.

### Effects of dopaminergic neurodegeneration and chronic treatment with pramipexole on behavioural flexibility

When considering the entire rat population with no distinction among FLEX, INT and INFLEX groups, there was a significant effect of the lesion (F_1, 153_ = 48.88, p<0.0001), time (F_2, 153_ = 60.75, p<0.0001) and lesion X time interaction (F_2, 153_ = 12.75, p<0.0001, **Fig 3A**). Viral-mediated expression of _α_-synuclein had no significant effect on inflexibility score when compared with baseline performances, but lesioned rats displayed worse performances than their sham counterparts after AAV surgery (p<0.001). Chronic treatment with PPX induced a significant worsening of behavioural flexibility in both sham and lesioned animals compared with post-AAV (p<0.0001, **Fig 3A**). However, this deleterious effect of PPX was more pronounced in lesioned than in sham animals (p<0.0001, **Fig 3A**).

**Figure 3:**
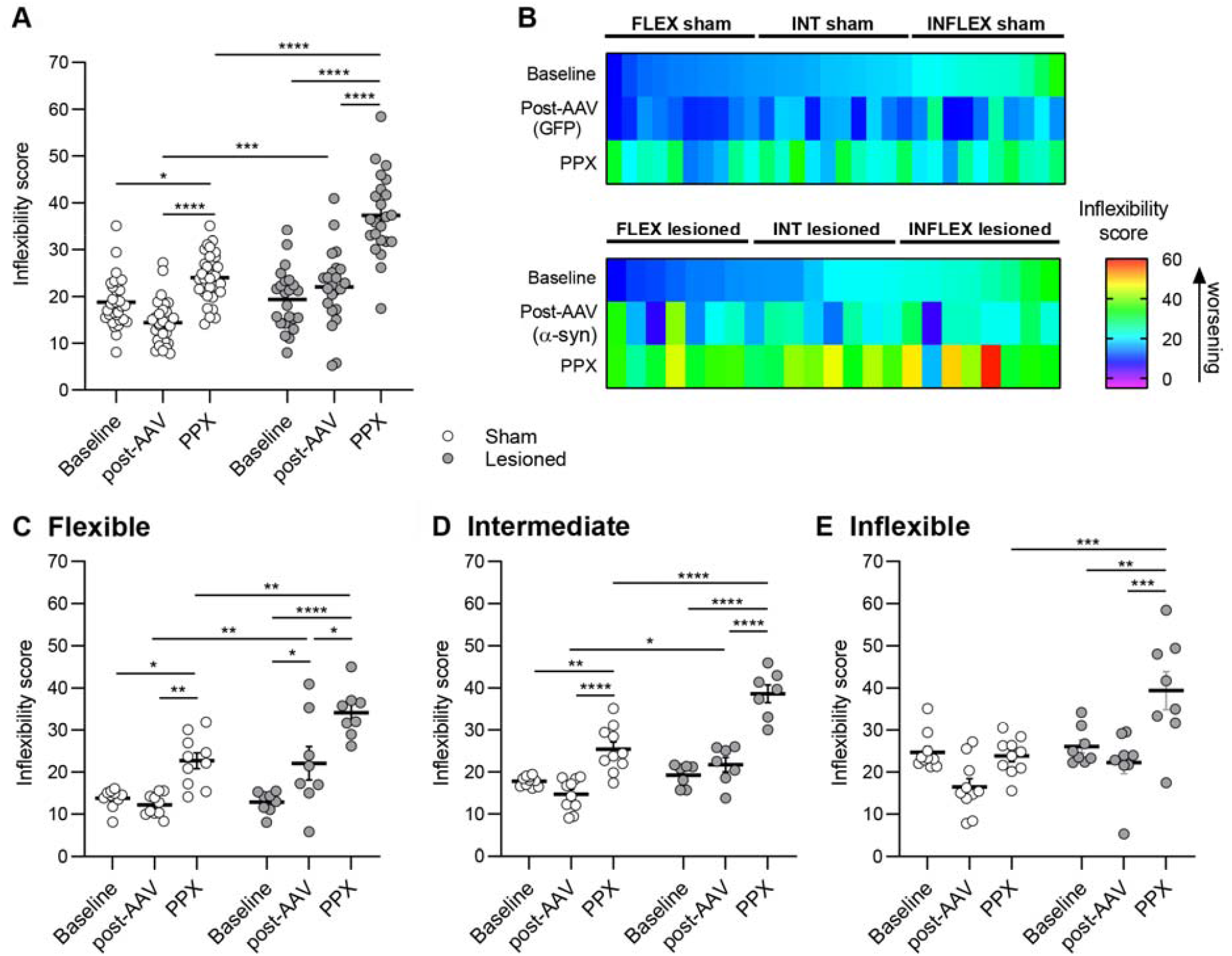
Effects of dopaminergic neurodegeneration and pramipexole on behavioural flexibility. **A**. Inflexibility score of whole sham and lesioned populations at Baseline, post-AAV injection and during chronic PPX treatment (PPX). **B**. Color plot illustrating the evolution of the inflexibility score for sham (GFP) and lesioned (_α_-syn) groups throughout the course of the experiment, each colored rectangle representing a single animal. Flexibility performances are represented at Baseline, post-AAV injection and following chronic PPX treatment (PPX): dark blue indicates a low inflexibility score and red a high inflexibility score. **C-E**. Inflexibility scores for sham *vs* lesioned group at Baseline, post-AAV injection and following chronic PPX treatment (PPX) for FLEX (**C**), INT (**D**) and INFLEX (**E**) groups.

When considering initial inflexibility score, this parameter was differentially affected both by the nigrostriatal lesion and by PPX (**Fig 3B**, individual animal’s inflexibility scores are represented by color-coded rectangles aligned vertically for each experimental time point). In FLEX animals there was a significant effect of the lesion (F_1, 48_ = 18.50, p<0.0001), time (F_2, 48_ = 32.68, p<0.0001) and lesion X time interaction (F_2, 48_ = 6.01, p<0.001, **Fig 3C**). Nigrostriatal lesion induced a significant deterioration of set-shifting compared with baseline (p<0.05). If PPX treatment worsened the inflexibility score both in sham (p<0.01) and lesioned FLEX rats (p<0.05), this deleterious effect was more pronounced in lesioned than in sham animals (p<0.01). In INT rats, there was a significant effect of the lesion (F_1, 45_ = 38.08, p<0.0001), time (F_2, 45_ = 60.50, p<0.0001) and lesion X time interaction (F_2, 45_ = 8.43, p<0.001, **Fig 3D**). Whereas baseline performances were not affected by the dopaminergic denervation, PPX induced an increase of the inflexibility score in both sham and lesioned rats (p<0.0001), and this deleterious effect was higher in lesioned than in sham rats (p<0.0001). In INFLEX rats, there was no difference between sham and lesioned rats at post-AAV, and only lesioned rats were affected by chronic PPX treatment (p<0.001 vs post-AAV and vs sham, **Fig 3E**).

These results indicate that _α_-synuclein induced nigral degeneration impairs behavioural flexibility only in FLEX animals and that the lesion potentiated the deleterious effect of PPX on behavioural flexibility in all animals. Moreover, the magnitude of the deleterious effect of PPX on flexibility negatively correlated with the inflexibility score before PPX treatment both in sham (r=-0.5841, p<0.001, **Fig 4A**) and lesioned animals (r=-0.4777, p<0.05, **Fig 4B**), indicating that in both groups the most flexible animals were the most impacted by PPX.

**Figure 4:**
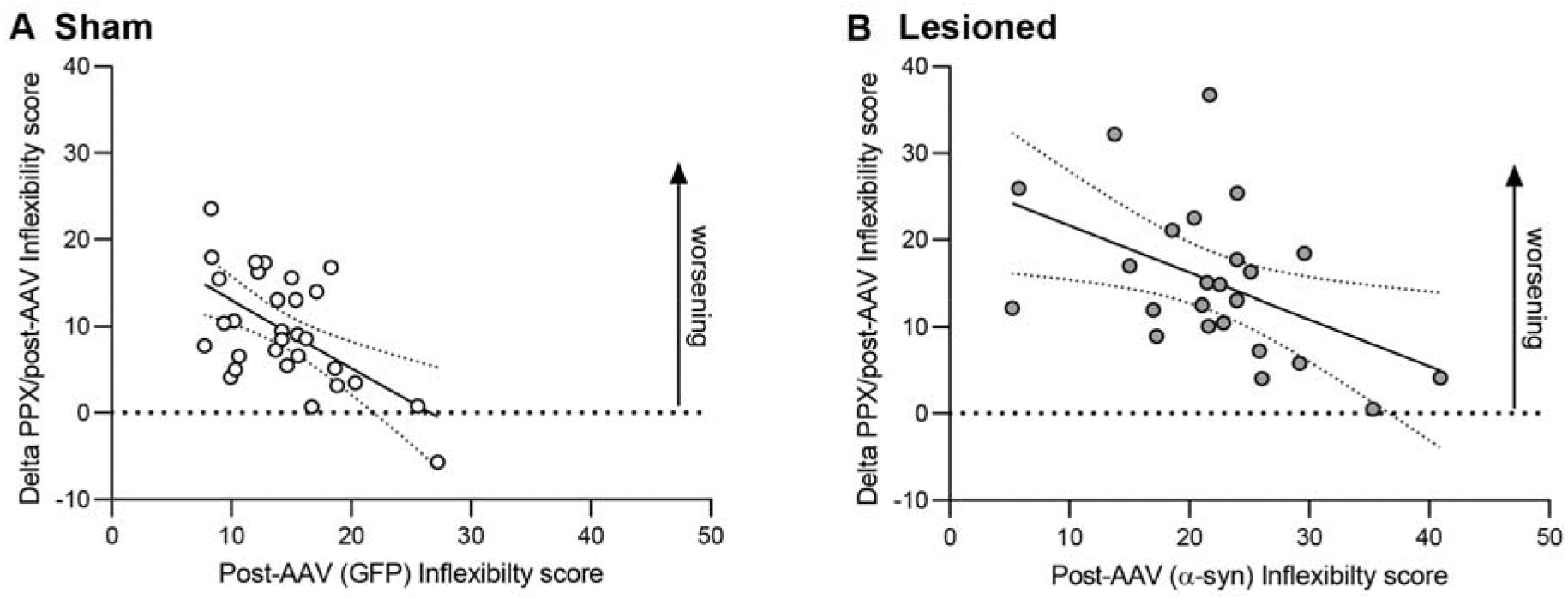
Relationship between the inflexibility score post-AAV injection and following chronic treatment with PPX. **A, B**. Correlations between the inflexibility score measured post-AAV injection and following PPX chronic treatment in sham (r=-0.5841, p<0.001, **A**) and lesioned rats (r=-0.4777, p<0.05, **B**).

### Effects of dopaminergic neurodegeneration and PPX on each dimension of the set-shifting task

As the set-shifting task involves rules based on two distinct perceptual dimensions (visual and egocentric), we assessed how each dimension was affected by the lesion and by PPX. The visual error rate was not impaired by the dopaminergic lesion or by PPX in FLEX and INT rats (**Fig 5A, B**). However, in INFLEX rats, there was a significant effect of the lesion (F_1, 48_ = 5.25, p<0.05) as well as lesion X time interaction (F_2, 48_ = 7.18, p<0.01), and PPX increased the visual error rate in INFLEX lesioned animals compared to their sham counterparts (p<0.001, **Fig 5C**). Regarding the egocentric dimension, there was a significant effect of the lesion (F_1, 48_ = 9.64, p<0.01) and time (F_2, 48_ = 40.83, p<0.0001) in FLEX animals that displayed a significant deficit after PPX compared to post-AAV (sham: p<0.001, lesioned: p<0.0001, **Fig 5D**). In addition, this deficit induced by PPX was greater in lesioned that in sham rats (p<0.01 **Fig 5D**). The error rates of INT sham and lesioned animals during the egocentric rule were also significantly higher after PPX treatment compared with post-AAV (p<0.01, **Fig 5E**). Conversely, the performances of INFLEX sham and lesioned animals in the egocentric dimension were unaffected by the lesion or by PPX (**Fig 5F**).

**Figure 5.**
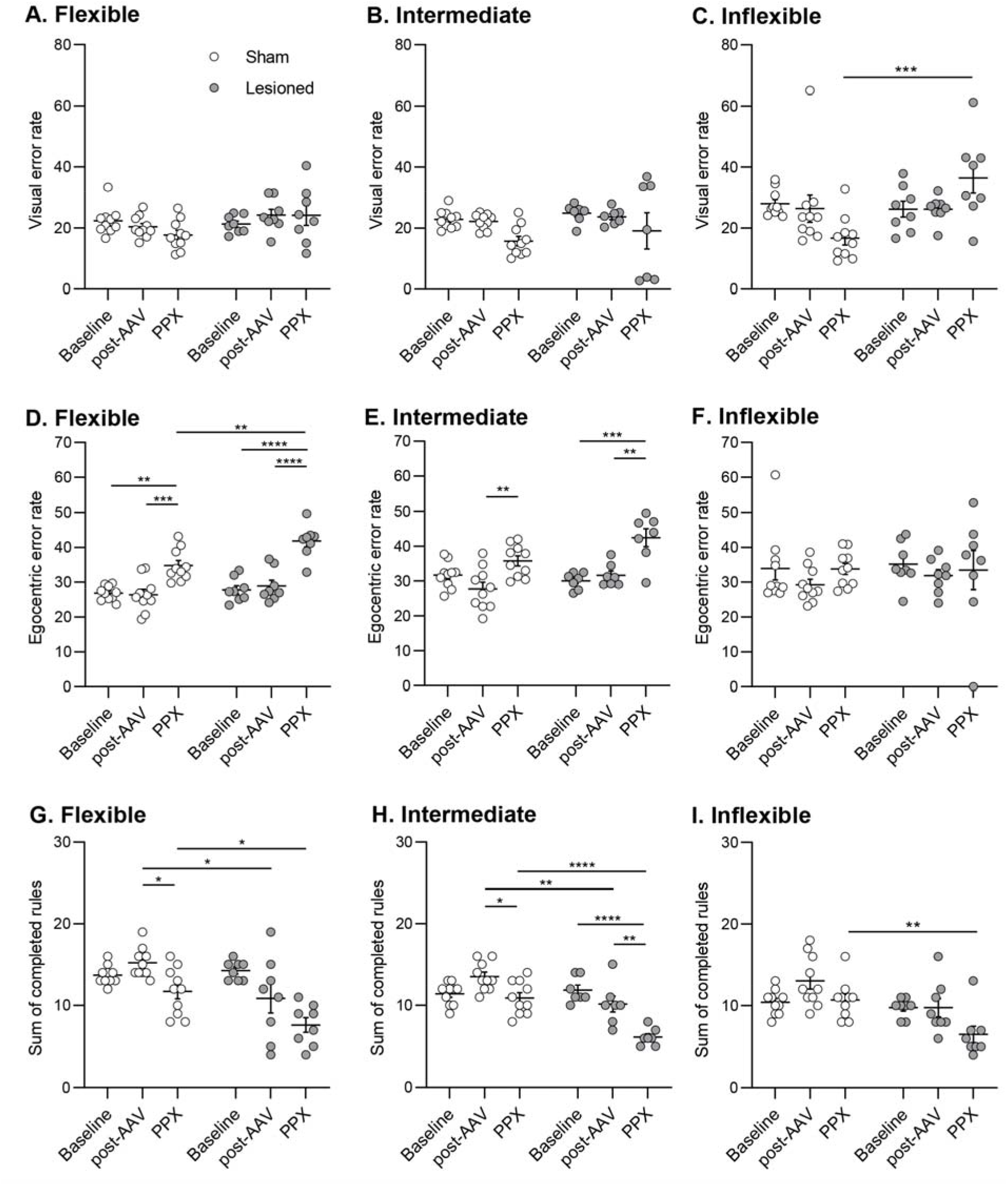
Effects of dopaminergic neurodegeneration and pramipexole on visual and egocentric error rates, and on the number of completed rules. **A-C**. Error rate in the visual rule at Baseline, after AAV injection (post-AAV), and following chronic PPX treatment (PPX) for sham and lesioned animals for FLEX (**A**), INT (**B**) and INFLEX (**C**) groups. **D-F**. Error rate in egocentric rule at Baseline, after AAV injection (post-AAV), and following chronic PPX treatment (PPX) for sham and lesioned animals for FLEX (**D**), INT (**E**) and INFLEX (**F**) groups. **G-I**. Number of completed rules at Baseline, after AAV injection (post-AAV), and following chronic PPX treatment (PPX) for sham and lesioned animals for FLEX (**G**), INT (**H**) and INFLEX (**I**) groups. *, p< 0.05 ; **, p<0.01 ; ***, p<0.001 ; ****, p<0.0001.

We also assessed the effect of the dopaminergic lesion and PPX on the number of completed rules during the task. In the FLEX group, there was a significant effect of the lesion (F_1, 48_ = 12.59, p<0.001), time (F_2, 48_ = 12.61, p<0.001) and lesion X time interaction (F_2, 48_ = 4.16, p<0.05). FLEX lesioned rats displayed a smaller number of completed rules at post-AAV compared with their sham counterparts (p<0.05, **Fig 5G**) that further decreased after PPX (p<0.05). A similar effect was observed in INT rats (lesion: F_1, 45_ = 25.8, p<0.0001, time: F_2, 45_ = 0.0001 and lesion X time: F_2, 45_ = 9.6, p<0.001), with lesioned rats displaying less completed rules post-AAV compared with shams (p<0.01, **Fig 5H**). The number of completed rules further decreased after PPX (sham: p<0.05, lesioned: p<0.01, lesioned vs sham: p<0.0001, **Fig 5H**). In INFLEX rats, the main effect of the lesion (F_1, 48_ = 16.12, p<0.001) and time (F_2, 48_ = 5.68, p<0.01) were evidenced only as a decreased number of completed rules after PPX in lesioned rats (p<0.01, **Fig 5I**).

## DISCUSSION

Cognitive impairment is a frequent non-motor feature of PD affecting up to 30% of newly diagnosed PD patients, ^21^ and can manifest under different forms including executive dysfunction such as cognitive and behavioural flexibility deficits.^2^ Although the pathophysiology of these non-motor symptoms is still incompletely understood, executive dysfunction is at least partially associated with a dopamine-dependent dysfunction of fronto-striatal circuits.^2^ Interestingly, altered activity of fronto-striatal circuits, decreased cognitive flexibility and set-shifting have also been reported in PD patients with ICD, ^12,13,21,22^ suggesting that interindividual differences in executive function prior to treatment initiation could contribute to the increased vulnerability of some patients to develop ICD.

This study aimed at investigating the contribution of nigrostriatal degeneration and dopamine replacement therapy with PPX on interindividual differences in behavioural flexibility. Using a validated set-shifting task ^23,24^ performed at the premorbid stage, after nigrostriatal degeneration and following chronic treatment with PPX in a rat model of PD, we demonstrate that nigrostriatal dopaminergic loss unequally affects behavioural flexibility depending on baseline performances. We further show that the deleterious effects of PPX on behavioural flexibility are increased in rats with nigrostriatal degeneration.

Our results provide the first experimental evidence that nigrostriatal dopaminergic neurons contribute to set-shifting abilities and they demonstrate that nigrostriatal degeneration has differential implications for set-shifting abilities depending on baseline performances. Indeed, a deleterious effect of nigral neurodegeneration was only observed in FLEX rats, whereas the performances of INT and INFLEX animals were unaffected. Stereological counts of nigral neurons demonstrated a similar level of denervation between FLEX, INT and INFLEX rats, indicating that this effect could not be attributed to a different magnitude of the lesion. These results are in agreement with clinical studies showing that PD patients display cognitive flexibility deficits as shown by an impaired ability to ignore irrelevant/distracting information, increased perseveration and learned irrelevance. ^5,25^ Even though the precise mechanisms underlying this involvement of the nigrostriatal pathway on set-shifting remain to be elucidated, there is evidence pointing toward a role for interindividual differences in dopamine homeostasis, as suggested by studies showing associations between polymorphisms in the catechol-O-methyltransferase gene and behavioural flexibility, both in the general population ^26^ and in PD patients.^27^ An association between polymorphisms in the dopamine transporter gene, fronto-striatal activation and set-shifting performances in PD patients further supports a role for interindividual differences in dopamine homeostasis as a substrate underlying behavioural flexibility. ^28^

Frontostriatal loops are implicated in behavioural flexibility and can be modulated by dopaminergic pathways at multiple levels. This modulation has been mostly studied in the prefrontal cortex and ventral striatum in rodents, and local infusion of dopamine agonists or antagonists can alter different aspects of behavioural flexibility. In the prefrontal cortex, local infusion of D1 or D2 antagonists impair set-shifting whereas D1 or D2 agonists have no effect. ^29^ In the nucleus accumbens, blockade of D1 or increased stimulation of D2 receptors both impair set-shifting, suggesting that a critical balance between activation of D1 and D2 receptors is required for optimal performances.^30^ Here, we provide evidence for a contribution of D3 receptor activation by demonstrating that systemic administration of PPX decreases behavioural flexibility. Importantly, our results indicate that nigrostriatal degeneration acts as an aggravating factor since this deleterious effect of PPX was markedly exacerbated in lesioned rats. In non-lesioned rats, PPX was found to differentially undermine set-shifting abilities depending on basal performances, with FLEX and INT animals being impaired whereas INFLEX animals were unaffected. Conversely, all lesioned rats were impaired by PPX regardless of their basal flexibility status. However, INFLEX lesioned rats showed a significant impairment only in the visual dimension whereas FLEX and INT rats were impaired only in the egocentric dimension, indicating a decreased ability to ignore the irrelevant and distracting visual information in the task. This deleterious effect of a D3 preferring agonist on set shifting abilities in rats with nigrostriatal degeneration is in agreement with previous work showing that a D3 antagonist was able to improve performances in an attentional set-shifting task in parkinsonian monkeys.^31^ Our results showing a pro-inflexible effect of PPX are also in line with previous findings showing that D3 knock-out mice display better performances in a set-shifting task.^32^

Even though executive impairment is common in PD, with one third of patients displaying some degree of executive dysfunction at the time of diagnosis ^33^, there is a noticeable clinical heterogeneity that may reflect the contribution of genetic factors (as discussed above), specific patterns of neurodegeneration and disease progression, as well as premorbid interindividual differences in cognitive performances.^6^ In this regard, our preclinical results help understanding such clinical heterogeneity by demonstrating that nigrostriatal degeneration leads to contrasting outcomes on behavioural flexibility depending on premorbid performance levels. Therefore, individuals who were the most flexible prior to nigral neurodegeneration might experience a drastic alteration of their behavioural flexibility, whereas poorly flexible individuals were much less affected. These findings also have implications for the pathophysiology of ICD in PD since clinical studies have reported impairment in cognitive flexibility and set-shifting in ICD.^13,15^

## CONCLUSION

By demonstrating that a loss of nigrostriatal neurons leading to homogenous motor impairments in a rat model of PD unequally impairs behavioural flexibility depending on premorbid performances, this study highlights non-motor outcomes of nigrostriatal neurodegeneration and helps understanding the bases of inter-individual variability of cognitive impairments in PD. Our results with PPX further highlight that the deleterious effect of this drug is not only exacerbated by the loss of nigrostriatal neurons, but involves impairments in different dimensions of the task between flexible and inflexible individuals, suggesting that behavioural processes subserving flexibility are differentially altered by the drug depending on individual baseline performances. Taken together, our results suggest that the vulnerability to develop behavioural flexibility impairments upon nigrostriatal neurodegeneration and treatment initiation could be linked to pre-morbid interindividual differences.

## METHODS

### Subjects

All experiments were approved by the local ethical committee (Comethea Poitou-Charentes C2EA-84, study approval #11562-2017092810291542) and performed under the European Directive (2010/63/EU) on the protection of animals used for scientific purposes. Male Sprague Dawley rats (n=52, 175g, Janvier, France) were housed on a reversed 12h cycle. After seven days of habituation, they were food restricted to 15g per day one week before behavioural studies. Water was available *ad libitum*.

### Flexibility procedure

This procedure was performed in operant chambers (Med Associates, Fairfax, VT, USA) equipped with two identical levers, each surmounted by a light cue. Before assessment of set-shifting abilities, five consecutive 1-hour training sessions were performed to train the rats to press the levers and to collect the reward in the magazine. Behavioural flexibility was then assessed before surgery (baseline, 16 sessions), 9 weeks after nigrostriatal degeneration (post-AAV surgery, 16 sessions) and during PPX treatment (acute PPX: first four days of treatment and chronic PPX: four last days of treatment), with a set-shifting task requiring discriminating between two rules based on two distinct perceptual dimensions: visual and egocentric.^23,24,34^ Experiments were performed daily during 45 minutes sessions. The visual and egocentric dimensions alternated during test sessions. During the visual rule, the active lever was determined by the position of the light cue above it. During the egocentric rule, only one lever (right or left) was active, independently of the position of the light cue (**Fig 1A**). For each rule, the position of the light cue was randomly determined and could occur above the same lever only for two consecutive trials.

A good lever choice led to obtain a 45mg sucrose pellet reward (TestDiet, Saint-Louis, MO, USA). Otherwise, a sound was triggered, and the house-light was turned on for twenty seconds (time-out). Rules alternated after ten correct responses. Each dimension (visual and egocentric) was presented at most twice per session. For each session, the percentage of incorrect responses (error percentage) and the number of completed rules were quantified. An inflexibility score was determined for each rat as the mean error percentage minus the number of completed rules over the last four sessions. Rats were clustered in tertiles, based on their baseline inflexibility score: Flexible (FLEX, n=17), Intermediate (INT, n=17) and Inflexible (INFLEX, n=18).

### AAV-α-synuclein-mediated lesion

Upon completion of baseline testing, FLEX, INT and INFLEX rats were randomly assigned to the sham or lesioned group. Rats were anesthetized with isoflurane (2% at 1.5 l/min) and placed in a stereotaxic frame (Kopf Instruments®, Tujunga, CA, USA). Vitamin A was applied on the eyes to prevent corneal dryness and 1mL of saline solution was subcutaneous injected to prevent dehydration. Pre-surgical analgesia was performed with ketoprofen (10 mg/kg, i.p.) and local analgesia with 2% xylocaine gel after disinfection with vetedine®. Two bilateral injections were performed in the SNc (in millimeters from bregma and dura, AP : -5,1 and -5,6 ; ML : +/-2,2 and +/-2 ; DV : -8) with 1µL of AAV2 expressing human A53T _α_-synuclein (CMVie/SynP-synA53T-WPRE, 5.2 10^13^ gcp/mL) or AAV2 expressing GFP (green fluorescent protein, CMVie/SynP-GFPdegron-WPRE, 3.7 10^13^ gcp/mL) at 0.2µL/min as previously described. ^19,35^ Post-surgical analgesia was performed with ketoprofen (10 mg/kg/day, i.p.) the day after surgery. Akinesia was estimated twice a day during four consecutive days using the stepping test as previously described ^36^ at baseline, 4 and 9 weeks after surgery, and following chronic PPX treatment.

### Treatment

After completion of the post-AAV surgery assessment of behavioural flexibility, all animals received PPX (Sequoia Research Products, Berkshire, UK) prepared daily in sterile saline and administrated subcutaneously at a dose of 0,3mg/kg/day for 21 days. Behavioural flexibility was tested 45 minutes after PPX injection to avoid interference with the early decreased locomotion occurring during the first 30 minutes after injection.^37^

### Tissue processing and histopathological analysis

Brain collection was performed after the last flexibility session. Sodium pentobarbital (120mg/kg, i.p.) was injected prior to an intracardiac perfusion with 200 mL 0,9% NaCl followed by 200 mL ice-cold 4% paraformaldehyde. Overnight post-fixation in the same fixative and cryoprotection in H2O/20% sucrose at 4°C were realized. Brains were frozen in isopentane at -40°C and stored at -80°C. Serial 50 µm coronal free-floating sections were collected and stored at -20°C in a cryoprotectant solution. After 3 washes in Tris Buffer Saline 1X (TBS, 15 minutes each), sections were incubated 10 minutes in a peroxidases blocking solution (S2023, Agilent Technologies, Santa Clara, CA, USA). After 3 washes, an incubation of 90 minutes in a blocking solution containing 3% BSA (bovin serum albumin) and 0.3% Triton in TBS 1X was performed. Sections were then incubated 18 hours at 4°C with mouse anti-TH (1/5000, MAB318, Sigma-Aldrich, Saint Louis, MO, USA) in blocking solution. Sections were washed again 3 times in TBS 1X, and then incubated 1 hour at room temperature (RT) with EnVision HRP system anti-mouse (K400111-2, Agilent Technologies, Santa Clara, CA, USA) in blocking solution. Then, sections were washed again three times in TBS 1X and immunoreactions were revealed with DAB peroxidase substrate (K346811-2, Agilent Technologies, Santa Clara, CA, USA) was performed. Finally, sections were mounted on gelatin-coated slides and coverslipped with DePeX.

TH positive neurons were counted using the optical fractionator method on every 6^th^ section of the SNc as previously described.^35^ Systematic random sampling was performed with the Mercator Pro V6.5 (Explora Nova, La Rochelle, France) software coupled with a Leica 5500B microscope. Following delineation of the SNc with x5 objective, counting was done with x40 objective.

### Statistical analysis

Data are expressed as mean□±□Standard Error of the Mean and analyzed using GraphPad Prism 8 software. Normality was tested with the Kolmogorov-Smirnov test. Data having a Gaussian distribution were analyzed using two-way analysis of variance (ANOVA) followed by Tukey’s post-hoc multiple comparisons test, except when comparing only two groups where a Student’s t-test was performed. Kruskal-Wallis test followed by Dunn’s post-hoc test, or Mann-Whitney test were applied when data did not follow a normal distribution. The following variables were analyzed: lesion (sham vs lesioned), time (baseline, post-AAV, PPX) and flexibility status (FLEX, INT, INFLEX). For all analyzes, a *p* value□<□0.05 was considered significant.

## Acknowledgements

The Université de Poitiers and the Institut National de la Santé et de la Recherche Médicale provided infrastructural support. This work was supported by Fondation de France (grant #00086205), région Nouvelle Aquitaine CPER 2015-2020 and FEDER 2014-2020 programs. The authors thank Dr Erwan Bezard for useful comments and suggestions on the manuscript. This study has benefited from the facilities and expertise of PREBIOS core facility at the University of Poitiers. The funders had no role in study design, data collection and analysis, decision to publish, or preparation of the manuscript.

## Data availability

The data that support the findings of this study are available from the corresponding author upon reasonable request.

## Authors contributions

conceptualization: MBM, MS, POF. Data acquisition: MD, EB, MF, MBM, POF. Statistical analysis: MD, MBM, POF. Writing of the first draft: MD, MBM, POF. Manuscript review and editing: MD, EB, MF, MS, MBM, POF

## Financial Disclosures of all authors for the preceding 12 months

EB, MF: none.

MBM has received grant support from France Parkinson. MD has received grant support from France Parkinson.

MS has received grant support from the IRESP « IRESP-19-ADDICTIONS-20», Nouvelle-Aquitaine Regional council and CPER-FEDER.

POF has received grant support from France Parkinson, Nouvelle-Aquitaine Regional council and CPER-FEDER.

